# Test-retest reliability of auditory MMN measured with OPM-MEG

**DOI:** 10.1101/2024.12.10.627674

**Authors:** Laszlo Demko, Sandra Iglesias, Stephanie Mellor, Katja Brand, Alexandra Kalberer, Laura Köchli, Stephanie Marino, Noé Zimmermann, Jakob Heinzle, Klaas Enno Stephan

## Abstract

In this paper, we report results from an investigation of auditory mismatch responses as measured by magnetoencephalography (MEG) based on optically pumped magnetometers (OPM). Specifically, as part of a quality control study, we examined the reliability and validity of auditory mismatch negativity (MMN) recordings, obtained with a newly installed OPM-MEG system. Based on OPM-MEG data from 30 healthy volunteers, measured twice with an established auditory MMN paradigm with frequency deviants, we examined the following questions:

First, we focused on construct validity and examined whether OPM-MEG measurements of MMN responses (in terms of event-related fields, ERFs) were qualitatively comparable to previous MMN findings from studies using EEG or MEG based on superconducting quantum interference devices (SQUIDs). In particular, we examined whether significant MMN responses measured by OPM-MEG occurred in a comparable time window and showed a similar topography as in previous EEG/MEG studies of MMN. Second, we quantified test-retest reliability of MMN amplitude and latency over two separate measurement sessions.

The results of our analyses show that MMN responses recorded with OPM-MEG are in good agreement with previously reported MMN results in terms of timing and topography. Furthermore, the comparison of group-level MMN topographies and timeseries shows excellent consistency across the two measurement sessions. Our quantitative test-retest reliability analyses at the sensor level indicate good reliability for MMN amplitude, but poor reliability for MMN latency.

Overall, our findings suggest that OPM-MEG measurements of auditory MMN (i) are comparable to results from EEG and SQUID-based MEG and (ii) show good test-retest reliability for amplitude measures at the sensor level. Notably, these results were achieved in an “out of the box” state of the OPM-MEG system, shortly after installation and without further optimisation. The reason for the insufficient reliability for MMN latency we observed is currently under investigation and represents an important target for future improvements.

## Introduction

Magnetoencephalography (MEG) based on optically pumped magnetometers (OPM-MEG) is a novel method to measure brain activity non-invasively in humans. OPM-MEG offers several advantages over conventional MEG as well as EEG (Brookes et al., 2022). For example, OPM sensors can be positioned within helmets or caps that fit individual head size; this enables measurements across the lifespan and patient-friendly short setup times. In contrast to classical cryogenic MEG based on superconducting quantum interference devices (SQUIDs), the setup is mobile and can be combined with dynamic field nulling, allowing participants to move their head or even their entire body (Boto et al., 2018; Holmes et al., 2023). These advantages are particularly valuable for groups who have difficulties sitting still for extended periods, such as children or certain groups of patients.

While OPM-MEG has tremendous potential, the technology is in its infancy, and its properties – in particular test-retest reliability (i.e. the stability or consistency of measurements in time) and construct validity (i.e. the agreement with results obtained by established technologies) – need to be characterised prior to routine practical applications. In particular, test-retest reliability has received little attention so far. One exception is a study that assessed test-retest scenarios for connectivity measures of resting state data (Rier et al., 2023). By contrast, we are not aware of any study evaluating test-retest reliability of standard paradigms for investigating evoked responses in translational and clinical contexts. In this study, we focused on the auditory mismatch negativity (MMN), a brain response to tones with unexpected frequency and/or duration that was originally discovered in EEG research (Näätänen et al., 2007). EEG-based MMN and its MEG equivalent (also referred to as MMNm), are very widely used in basic and clinical neuroscience (Näätänen et al., 2007; Recasens and Uhlhaas, 2017). The psychometric properties of MMN, including test-retest reliability, have been examined for both SQUID-based MEG (Recasens and Uhlhaas, 2017) and EEG (e.g., Wang et al., 2021). While results vary, depending on the type of MMN paradigm used (e.g. frequency MMN or duration MMN) and the particular feature of the MMN studied (e.g. amplitude or latency), the overall conclusion from these studies is that the measures of MMN show good test-retest reliability, thus supporting its use for research and clinical applications.

In this paper, we report results from a quality control study that assessed the test-retest reliability and construct validity (i.e. consistency with previously published MMN findings) of MMN measurements obtained with a newly established OPM-MEG laboratory setup. It is worth noting that this quality control assessment was performed for an “out-of-the-box” OPM-MEG system, directly after its installation and without further optimisation, in a “plug and play” state. For example, OPM-MEG data were acquired in the absence of quantitative procedures for optimising helmet-brain realignment across subjects but relied on manual placement and checks of OPM helmet positions.

In this quality control study, we addressed the following two questions:

First, we focused on construct validity and examined whether we could obtain MMN responses (in terms of event-related fields, ERFs) with our OPM-MEG setup that were qualitatively comparable to previous MMN findings in the MEG (Maess et al., 2007; Recasens and Uhlhaas, 2017) and EEG (Weber et al., 2022) literature. In particular, we focused on the following qualitative comparisons: Do significant MMN responses measured by OPM-MEG

1. occur in a comparable time window as MMN responses measured by EEG under the same paradigm (Weber et al., 2022)?
2. occur in a comparable time window as MMN responses measured by SQUID-based MEG in previous studies that used different auditory MMN paradigms but also focused on frequency deviants (Maess et al., 2007)?
3. show a topography that is similar to the one in conventional MEG studies (Maess et al., 2007; Recasens and Uhlhaas, 2017)?

Second, we assessed the stability of the measurements by quantifying the test-retest reliability (Liljequist et al., 2019). This test-retest analysis can be compared to previous assessments of MMN test-retest reliability for SQUID-based MEG, for example as reported by (Recasens and Uhlhaas, 2017).

## Methods

All steps of data preprocessing and statistical analysis were pre-specified in an analysis plan ex ante. Prior to data analysis, this analysis plan was preregistered on ZENODO (Demko et al., 2024). Any deviations from the prespecified analysis are explained in the Methods and Results sections below.

### Participants

30 participants (16 female/14 male, age (mean±std): 29.7±7.8 years, range: 20-48 years) were recruited for this study. All participants gave their written informed consent. The study was approved by the ethics committee of the ETH Zürich (application number: 24 ETHICS-163). Each participant was invited to two separate sessions recorded at the same time of the day (± 2 hours). The second session was recorded 1-3 days after the first session. For each individual, we assured that the OPM helmet was placed in similar positions between the two sessions by adjusting the distance between the rim of the helmet and the tip of the participant’s nose in the second session to match the one measured in the first. The experiment was started within a few minutes after the helmet was placed.

### OPM-MEG data acquisition

Continuous OPM-MEG data were recorded inside a magnetically shielded room (MSR) optimized for OPM operation using demagnetization coils and a matrix coil system for dynamic field nulling (Cerca Magnetics Limited, Nottingham, UK). The OPM data of a 64-sensor array of triaxial zero-field OPM sensors (QuSpin Inc., Louisville, CO, USA) were sampled at 1200 Hz using a series of National Instruments (NI, Austin, TX, USA) NI-9205 16-bit analogue to digital converters interfaced with LabVIEW (NI, Austin, TX, USA). The sensors were mounted inside 3D-printed rigid helmets (Cerca Magnetics Limited, Nottingham, UK). For each participant we used the helmet that optimally fitted their head size. Participants were seated on an armchair in front of a screen within the MSR.

### Auditory MMN paradigm

During the experiment, participants listened to a tone sequence consisting of low (440 Hz) and high tones (528 Hz) produced by an Arduino microcontroller board (Arduino SA, Chiasso, Switzerland). Tones were delivered using E-A-RTONE™ 3A insert earphones (3M Auditory Systems, Indianapolis, IN, USA). The stimulus onset asynchrony (SOA) of tones was 570 ms with a stimulus duration (SOT) of 70 ms. A total of 1800 stimuli were presented using exactly the same sequence as in a previous study (Weber et al., 2022). The probability of hearing either the low or the high tone at any point during the sequence varied according to a pre-defined schedule (see Figure 1 in Weber et al. 2022). During the entire experiment, participants performed a visual distractor task, which required them to focus their attention on a square presented centrally on a screen in front of them and to indicate via a corresponding button press whenever the square opened either on the left or the right side. For a detailed description, please see (Weber et al., 2022).

**Figure 1:**
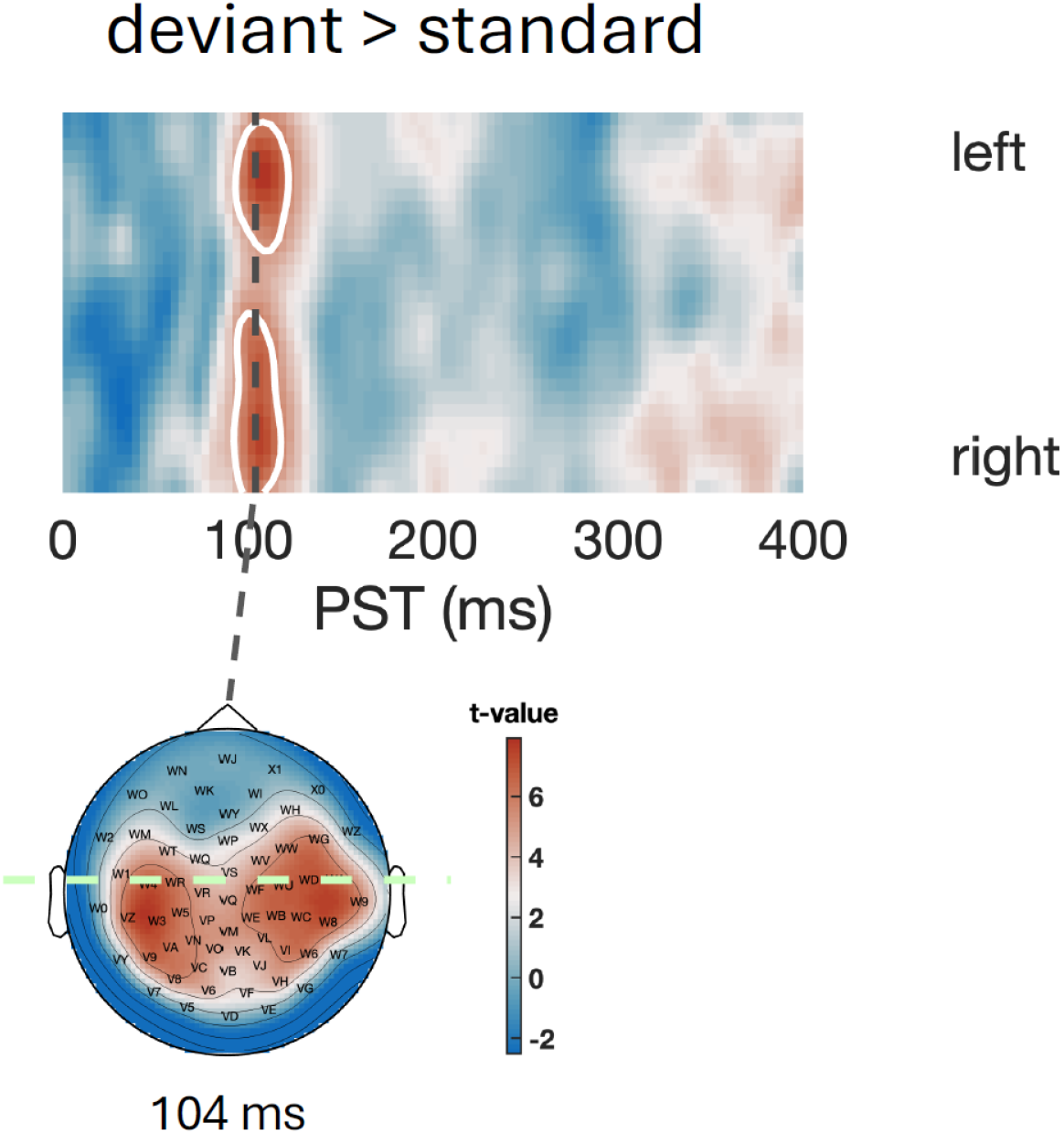
Group level difference of ERF magnitude between deviant and standard trials. Top: T-values (see colorbar) are shown for a section (left to right through the maximum) as indicated below. White contours indicate the two significant clusters. Bottom: Topography of t-map at time 104 ms showing both peaks.

### Preprocessing of OPM-MEG data

The raw OPM-MEG data were preprocessed using MATLAB and SPM12. The following steps were performed sequentially on the continuous raw MEG recordings: First, the power spectral density (PSD) was calculated for all channels to check the noise floor and identify bad channels. Channels with PSD values outside the range [7, 200] *fT*/√*Hz* within the [65, 75] *Hz* frequency range were excluded from further analysis. The data were then high-pass filtered using a fourth-order Butterworth filter with cutoff frequency 0.1 Hz, low-pass filtered using a fourth-order Butterworth filter with cutoff frequency 30 Hz.

Next, we employed adaptive multipole models (AMM; Tierney et al., 2024) to remove global signals not originating from the brain. The parameters of AMM were set to use internal and external harmonic order of 10 and 5, respectively, with a correlation limit of 0.98.

For the event-related analysis, the data were epoched into 500 ms segments around tone onsets, using a pre-stimulus time window of 100 ms. At this point, we checked the data quality of individual trials. Trials in which the standard deviation of the signal at any one channel was larger than three times the average standard deviation of the signal at that channel across all trials were labeled as bad trials and excluded from further analysis. Finally, we downsampled the data to 240 Hz.

In order to facilitate the comparison of our findings with previous MMN results from the EEG literature, the settings of the high-pass filter (0.1 Hz) were chosen in accordance with (Weber et al., 2022) and more general recommendations for high-pass filtering for EEG data (Tanner et al., 2015). However, it is unclear to what degree these recommendations apply to MEG, and higher cutoffs are chosen in many EEG/MEG studies of evoked responses. As a control and sensitivity analysis, we therefore re-ran the entire analysis pipeline using a high-pass filter of 1 Hz, as prespecified in our analysis plan.

### Trial definition

For the analysis, we defined deviants and standards within the stimulus sequence in the same way as in a previous study that used the same oddball MMN design we use here (Weber et al., 2022). A deviant trial was defined as the first auditory stimulus of a different frequency after at least 5 repetitions of the same tone. A standard trial was defined as the sixth repetition of the same stimulus type. This procedure yielded 106 standard and 119 deviant trials from a total of 1800 trials. It is worth noting that our analysis did not consider the factor ‘volatility’, i.e. the change between stable and instable phases, which was also assessed in the previous study (Weber et al., 2022).

### Reason for exclusion of trials and data sets

In order to assure high quality of the data that entered the statistical analysis, a set of criteria for exclusion of datasets or individual trials were defined a priori (see the Analysis Plan; Demko et al., 2024). Reasons for excluding data sets from the analysis were:

- Technical issues of data acquisition that were not visible during the measurements but only become apparent during quality control and preprocessing of the data.
- Excessive number (more than 20%) of bad trials as labeled in the previous step or more than 10% of bad channels.

### Realignment of helmet orientation

As mentioned above, the current analyses were performed without any procedures for optimising helmet-brain realignment across subjects. In the analysis plan (section 5.3), we had expressed the intention of attempting a post-acquisition realignment procedure but later realised that such a procedure is more challenging than anticipated and requires further development and validation. We therefore relied on manual placement and checks of OPM helmet positions, without any post-acquisition realignment.

### Computation of event related fields and magnetic field magnitude

Next, for each participant, we computed the ERF for deviant and standard trials, respectively. This was achieved by averaging the vector in each sensor over all corresponding trials and applying a baseline correction, i.e. subtraction of the average of the 100 ms pre-stimulus time-window for each channel (i.e. sensor direction). Note that this is equivalent to subtracting the average (between -100 ms and 0 ms) magnetic field vector of each three-dimensional sensor trace.

Here, we deviated slightly from the analysis plan and computed the averages manually, instead of using a general linear model (GLM) in SPM. As a consequence, the conversion of sensor data into images was performed later, just before the group level analysis (see below). All further statistical analyses of these images did rely on GLMs in SPM12, as prespecified.

In order to reduce the signal from the tri-axial OPM sensors (a three-dimensional vector) into a single value for statistical analysis by the GLM, we computed the magnetic field magnitude at each sensor location as the length of the magnetic field vector using the L2 norm. This approach is not a standard choice in classical MEG research but exploits the fact that our new setup contains the most recent generation of OPM sensors, i.e. tri-axial sensors. In order to compare the mismatch ERFs with previous studies using conventional SQUID-based MEG, we repeated all analyses using the z-component of the sensors which reflects the radial component of the magnetic field perpendicular to the scalp, similar to the SQUIDs in a conventional MEG. Note that the value of the z-component can also be negative, but does of course contribute to the field magnitude.

### Conversion to images and smoothing

For the group analysis, the data containing the magnitude (or z-component) of the two conditions were converted into 3D (scalp (2D) × within trial time points) images. This resulted in two images (one per condition) per participant. Subsequently, spatial smoothing was performed with a Gaussian kernel of 16 mm FWHM in both spatial dimensions. It is important to note that smoothing supports the group level analysis by making it less sensitive to potential variations of sensor placement across subjects. As mentioned above, the realignment of the two sessions within each subject was carefully controlled by adjusting the distance between the nose and the rim of the helmet to exactly match the first session.

### Group level analysis

Using the L2 norm based magnitude images of the average ERF, we examined the average positive MMN (difference between deviant and standard trials) at the group level. This was implemented as a one-tailed paired t-test. Note that there is a clear hypothesis regarding the direction of the effect: the magnitude of the ERF should be increased for deviant trials.

For the radial (z) component of the ERF, we also assessed the difference between deviant and standard but used two-tailed paired t-tests. This reflects the fact that the z-component reflects the direction of the field along one dimension and stronger activation during deviants can result in a positive or negative effect, depending on the direction of the field. In SPM, we tested therefore separately for positive and negative effects of mismatch and applied Bonferroni correction for two tests, i.e. a threshold of p<0.025 per test.

For all group level analyses, we applied a significance threshold of p<0.05, family-wise error (FWE) corrected for multiple comparison at the peak level using Gaussian random field theory. Notably, these group analyses were conducted for both sessions separately and the results are compared qualitatively by displaying the resulting clusters and t-maps.

### Test-retest reliability

In order to assess the test-retest reliability of the MMN measured with the new OPM setup, we conducted the following analysis using a strategy similar to (Recasens and Uhlhaas, 2017).

We used the group results of the first session to select the sensors associated with the significant activations of the group analysis. Specifically, we selected all sensors that belonged to (i) the global maximum peak and (ii) the first large cluster starting at around 100 ms, respectively, separately in the left and right hemisphere. This resulted in two sensors or sets of sensors, respectively, one for each hemisphere. We then computed the average timeseries, separately for the left and right sensors. Based on these timeseries, we extracted features of interest for assessing test-retest reliability, similar to (Recasens and Uhlhaas, 2017). Specifically, we extracted the following two features for each subject, each of the selected sensor groups and each session:

1. Average amplitude between -10ms to +10ms around the maximum (minimum) which, as described above, was determined within a time window of ±50 ms around the group maximum (minimum).
2. Response latency (in ms after stimulus onset) of the maximum (minimum) which was determined within a time window of ±50 ms around the latency of the group effect.

Maximum or minimum (for the radial component only) were chosen depending on whether the group analysis revealed a maximum or minimum.

Using these measures, we computed the intra-class correlation coefficient ICC(3,1) according to the definition by Shrout & Fleiss (1979) and as reviewed in (Liljequist et al., 2019) as ICC(C,1):

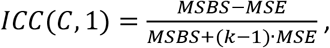

where k=2 is the number of measurements, and the mean square between subjects (MSBS) and mean squared error (MSE) are computed as in (Liljequist et al., 2019). Confidence intervals for the ICC were computed according to (McGraw and Wong, 1996) using the Matlab ICC toolbox (Salarian, 2024).

## Results

In the following, we first briefly present some general aspects of data quality. Then, we report group-level MMN analyses and test-retest reliability results for magnetic field strength (L2 norm). Subsequently, we present results based only on the radial (z) component of the magnetic field which is comparable to SQUID-based signals in a conventional MEG and thus allows for comparing the resulting topographies and traces to previous MEG studies of the MMN.

### Data quality

From the 30 participants measured, all but two had OPM-MEG data with sufficient quality to perform the planned analysis. In these two participants, more than 20% for the trials were labeled as “bad” (48 and 58 out of the 225 trials (standard and deviants), respectively). For the 28 participants who entered the analysis, we excluded on average 3.5±2.0 (mean±std; range 0 - 8) channels and 4.9±5.3 (mean±std; range 0 - 20) trials.

### Magnetic field magnitude (L2 norm) - Group level analysis

The comparison between L2 norm based magnetic field amplitudes in deviant and standard trials resulted in significant bilateral activation (p<0.05, FWE corrected for multiple comparison at the peak level). Specifically, there were two significant clusters both with a peak at 104 ms, one in the right hemisphere (SPM coordinates: x = 51 mm, y = -19 mm; T = 7.57, P_FWE-corr_ < 0.001, cluster size: 693 vox.) and one in the left hemisphere (x = -38, y = -25 mm; T = 8.02, P_FWE-corr_ < 0.001, cluster size: 565 vox.). See Figure 1 for an illustration of the statistical parametric map of the event related field. The sensor positions closest to the two peaks were WD (right) and W3 (left), respectively.

To compare the consistency of group level results between the two sessions, we performed exactly the same analysis for the second experimental session. The same two prominent peaks were observed in session 2. They occurred at 117 ms (x = 60 mm, y = -36 mm; T = 7.28, p_FWE-corr_ = 0.002) in the right hemisphere and at 108 ms (x = -42 mm, y = -25 mm; T = 8.66, p_FWE-corr_ < 0.001) in the left hemisphere. Note that in the second session there was one large cluster extending over both hemispheres (cluster size: 1363 vox.). There were two additional activations, one small cluster at 125 ms (x = 21 mm, y = 24 mm; T = 4.80, p_FWE-corr_ < 0.012, cluster size: 10 vox.) and one single voxel (x = -34 mm, y = -36 mm; T = 4.47, p_FWE-corr_ < 0.043). Figure 2 illustrates the similarity of the time courses and topographies of the event related fields in the two sessions. The visual similarity of the activations across the two sessions is striking and indicates very good reliability of group results.

**Figure 2:**
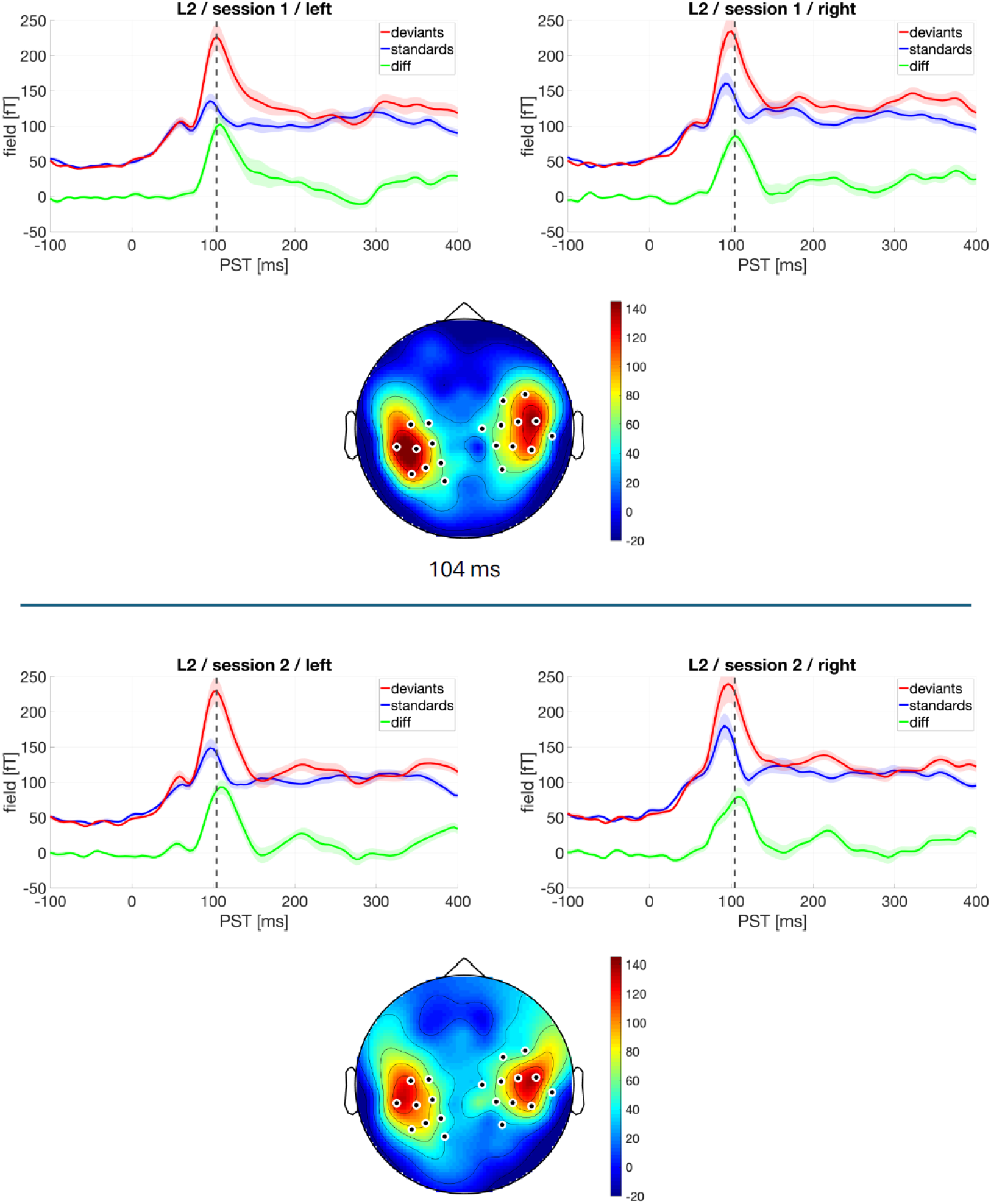
Comparison of group level results across sessions – magnetic field amplitude (L2 norm). Average traces of left and right hemisphere sensors are displayed for deviants (red), standards (blue) and their difference (green, MMN). The shaded area illustrates that standard error of the mean. The selected sensors are indicated by the white stars. Note that the topographical plot is shown for the timing of the first session and that the sensors were selected based on data from the first session only.

### Magnetic field magnitude (L2 norm) – test-retest reliability analysis

After having demonstrated the reproducibility of group level results, we tested the within-subject consistency in a test-retest reliability analysis. For this, we selected the sensors that belonged to (i) the global peak activation and (ii) the first large cluster starting at around 100 ms, respectively, and computed the average time courses separately in the left and right hemisphere. Next, we extracted the maximum (amplitude and timing) in a window of ± 50ms around the group peak for each individual and session. Based on these values, we computed the ICC for amplitude and peak time, respectively. Concerning (i), the sensors showing maximum MMN responses in the left and right hemisphere, respectively, were located at the following coordinates: x = 51 mm, y = -19 mm (right) and x = -38, y = -25 mm (left). Their timeseries showed good test-retest reliability for amplitude: ICC = 0.80 (right hemisphere) and ICC = 0.74 (left hemisphere). By contrast, test-retest reliability was poor for latency measures: ICC = 0.43 (right hemisphere) and ICC = 0.35 (left hemisphere). Table 1 summarizes all ICC values including confidence intervals and statistics. . Concerning (ii), this resulted in the selection of 11 sensors in the right and 9 sensors in the left hemisphere. Amplitude measures showed good test-retest reliability: ICC = 0.79 (right hemisphere) and ICC = 0.77 (left hemisphere). By contrast, latency measures (timing of the peak) showed very poor test-retest reliability: ICC = 0.34 (right hemisphere) and ICC = -0.20 (left hemisphere). See Table 1 for a summary of ICC values and their confidence intervals.

**Table 1:** ICC values based on magnetic field magnitude. ICC values for amplitude and latency are given for both methods. 95% confidence intervals were computed according to (McGraw and Wong, 1996) and p-values result from a comparison against the null hypothesis ICC = 0.

| sensor selection | measure | hemisph. | ICC(C,1) | 95%-CI | p-value |
| --- | --- | --- | --- | --- | --- |
| peak | amplitude | left | 0.74 | 0.51 - 0.87 | <0.001 |
|  |  | right | 0.80 | 0.61 - 0.90 | <0.001 |
|  | latency | left | 0.35 | -0.01 - 0.64 | 0.030 |
|  |  | right | 0.43 | 0.07 - 0.69 | 0.010 |
| cluster | amplitude | left | 0.77 | 0.56 - 0.88 | <0.001 |
|  |  | right | 0.79 | 0.60 - 0.90 | <0.001 |
|  | latency | left | -0.20 | -0.53 - 0.18 | 0.854 |
|  |  | right | 0.34 | -0.03 - 0.63 | 0.036 |

### Radial field (z) component – Group level analysis

In order to establish a closer comparison to previous MEG studies of auditory mismatch activity, we repeated the above analyses but using only the radial component of the magnetic field which is perpendicular to the skull. This corresponds to the z-direction of the sensors in the OPM helmet arrangement. The ERF differences between deviant and standard were assessed using a paired two-tailed t-test, which was implemented in SPM by two separate t-test at p<0.025. This allows for both positive and negative deviations to be detected. As for the magnitude, we observed a symmetric, bilateral MMN with peak times at 113 ms (left) and 146 ms (left). However, the clusters extended much longer in time from roughly 90 ms to 190 ms. Figure 3 shows the statistical maps of this analysis. A list of all significant clusters is provided in Table 1. The sensors closest to these peaks were WD (right) and W4 (left).

**Figure 3:**
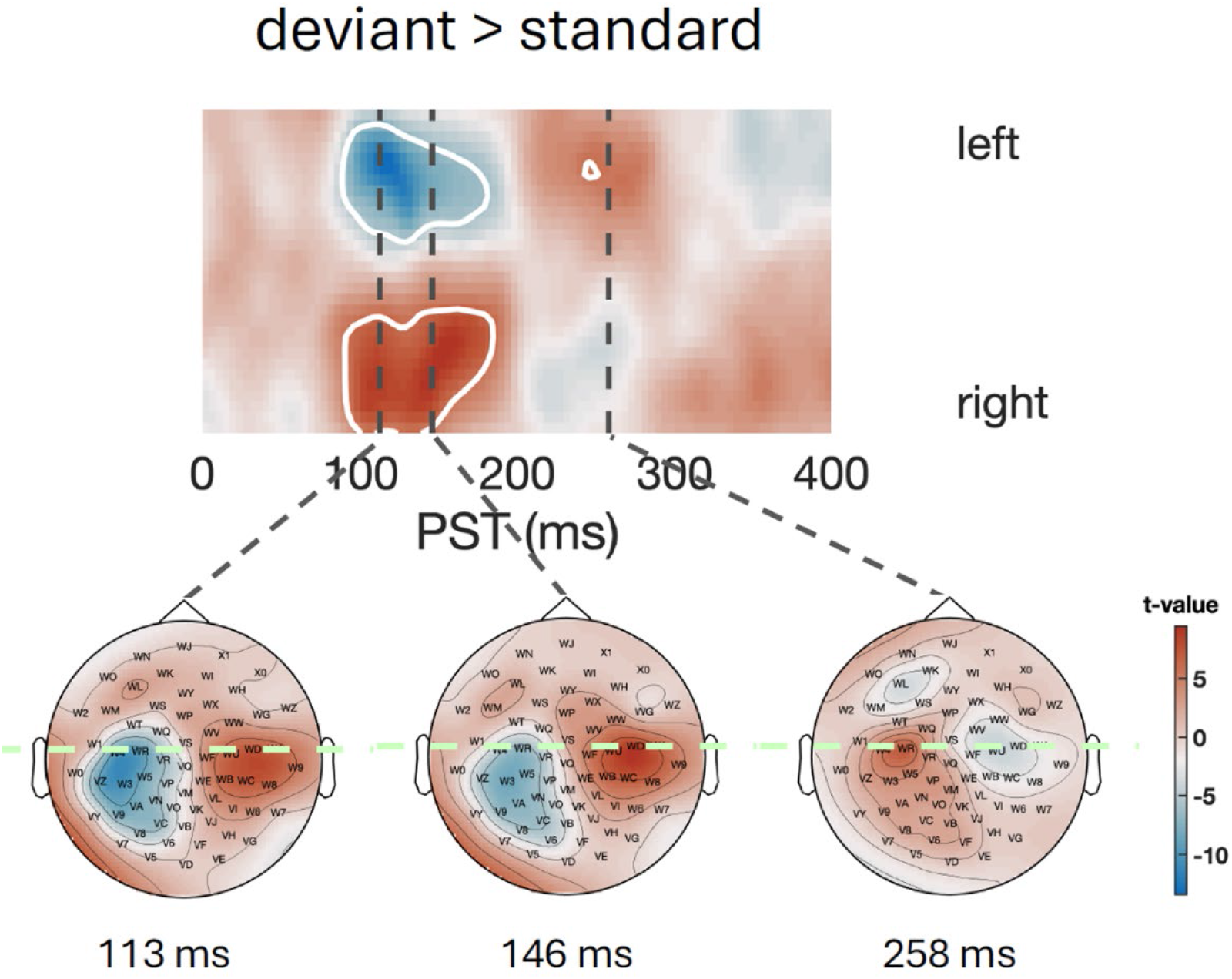
Group level difference of radial field component between deviant and standard trials. Top: T-values (see colorbar) are shown for a section (left to right through the maximum) as indicated below. White contours indicate the two significant clusters. Bottom: Topography of t-map at times 113 ms, 146 ms and 258 ms.

The second session showed highly similar results. Again, there were two main clusters that showed a temporally extended MMN lasting for nearly 100 ms (see Table 2).

**Table 2:** Summary of statistical results for the analysis of the radial component. All significant clusters are listed for the two sessions. Cluster size is given as the number of above threshold ‘voxels’ in the 3D (2D-space x time) spm image. Positive t-values indicate that the difference of the radial component between deviants and standards is positive, while negative t-values indicate that it is negative. Note that a negative t-value also occurs when the magnetic field is stronger in the negative direction, still indicating a stronger response for the deviant. For an interpretation of these values, please consider Figure 4, where the values of the magnetic field are shown. * These values are given for the peak of the cluster.

|  | time* | hemisph. | location*<br>(x,y) in mm | size<br>(voxels) | T-value* | p<sub>FWE-corr</sub>* |
| --- | --- | --- | --- | --- | --- | --- |
| **Session 1** | 146 ms | right | (42,-14) | 1534 | 9.53 | <0.001 |
|  | 113 ms | left | (-38,-19) | 1998 | -13.39 | <0.001 |
|  | 258 ms | left | (-26,-3) | 63 | 7.28 | 0.002 |
|  | 100 ms | left | (-38,50) | 14 | 7.41 | 0.002 |
|  | 96 ms | right | (60,29) | 11 | -7.67 | <0.001 |
| **Session 2** | 125 ms | right | (47,-19) | 670 | 8.77 | <0.001 |
|  | 121 ms | left | (-38,-30) | 1884 | -11.74 | <0.001 |
|  | 213 ms | left | (-30,-30) | 2 | 5.69 | 0.049 |
|  | 396 ms | right | (-34,-19) | 1 | -5.73 | 0.044 |

Figure 4 compares the topographies and traces of the radial magnetic field for the two sessions. Note that the mismatch activity in this case is extended in time, lasting from around 100 ms to 200 ms after stimulus onset. This is in line with the duration of the MMN in the EEG study of Weber et al. (2022). The topographies display a classical MMN layout as observed e.g. in (Maess et al., 2007; Recasens and Uhlhaas, 2017). As for the L2 norm based magnetic field magnitude, the group results from the second session are highly similar, indicating high reliability of group-level results.

**Figure 4:**
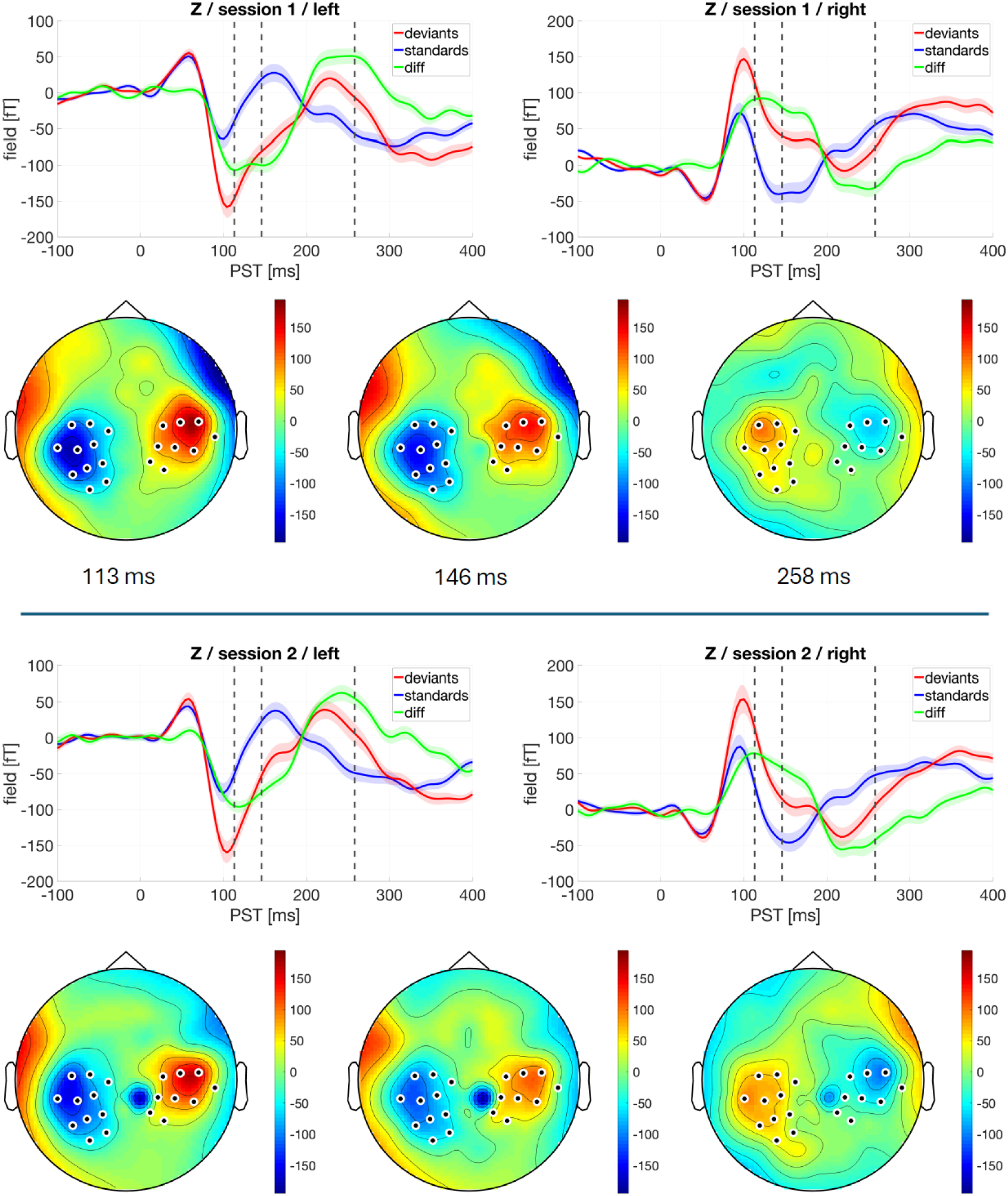
Comparison of group level results – radial magnetic field. Average traces of left and right hemisphere sensors are displayed for deviants (red), standards (blue) and difference (green, MMN). The selected sensors are indicated by the white stars. Note that the topoplot is shown for the timing of the first session and that the sensors were selected based on data from the first session only.

### Radial field (z) component - test-retest reliability analysis

For the test-retest analysis of the radial field, we followed the same strategy as for the magnitude. Again, we first considered the traces of the sensors closest to the minimum (left) and maximum (right) of the t-statistics. Next, we averaged the traces of the 9 sensors within the right cluster and the 11 sensors within the left cluster. For both peak and cluster analysis amplitude showed a test-retest reliability above 0.5, but was not as reliable as for the magnetic field amplitude. Latency showed again poor reliability clearly below 0.5. See Table 3 for a summary of ICCs based on the radial field component.

**Table 3:**
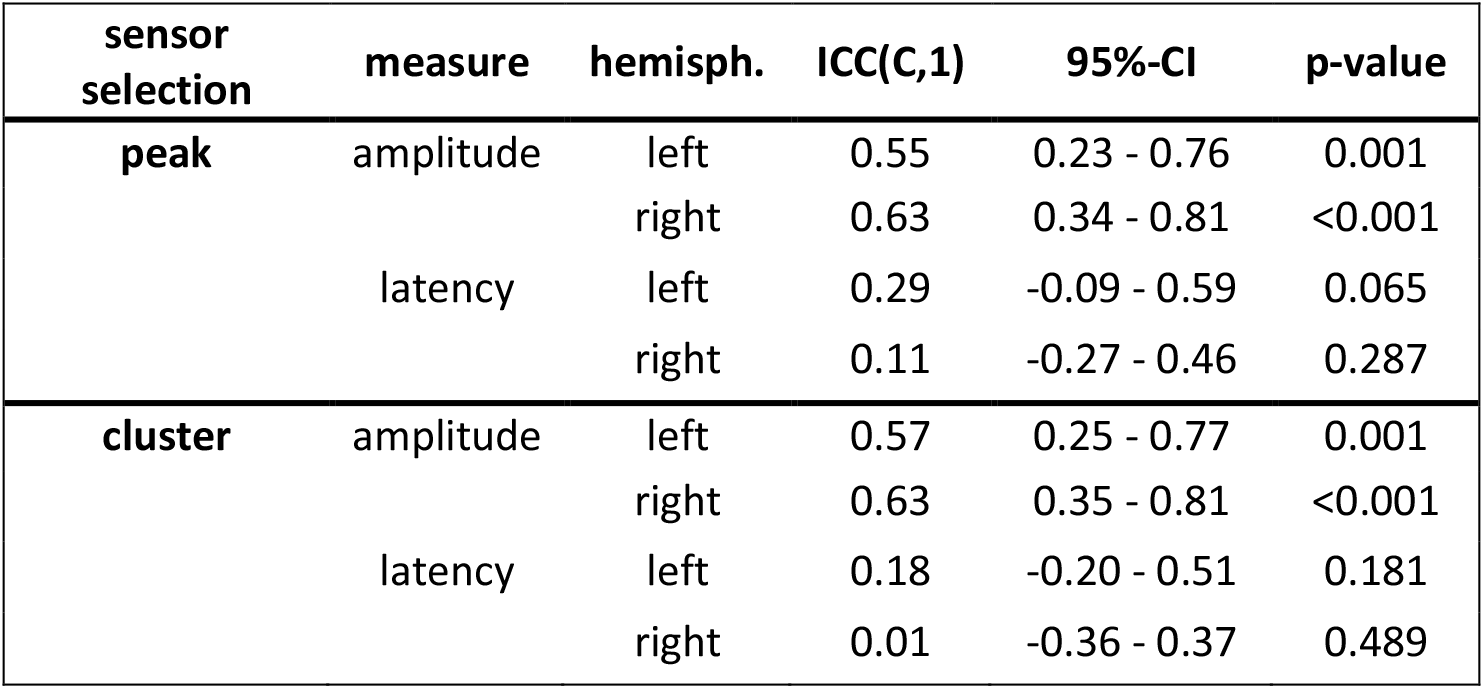
ICC values for analysis based on radial field component. ICC values for amplitude and latency are given for both the peak sensor as well as the average trace of sensors within the significant cluster. 95% confidence intervals were computed according to (McGraw and Wong, 1996) and p-values result from a comparison against the null hypothesis ICC = 0.

| sensor selection | measure | hemisph. | ICC(C,1) | 95%-CI | p-value |
| --- | --- | --- | --- | --- | --- |
| peak | amplitude | left | 0.55 | 0.23 - 0.76 | 0.001 |
|  |  | right | 0.63 | 0.34 - 0.81 | <0.001 |
|  | latency | left | 0.29 | -0.09 - 0.59 | 0.065 |
|  |  | right | 0.11 | -0.27 - 0.46 | 0.287 |
| cluster | amplitude | left | 0.57 | 0.25 - 0.77 | 0.001 |
|  |  | right | 0.63 | 0.35 - 0.81 | <0.001 |
|  | latency | left | 0.18 | -0.20 - 0.51 | 0.181 |
|  |  | right | 0.01 | -0.36 - 0.37 | 0.489 |

### Analyses with alternative cutoff for high-pass filtering

As described in the methods section and as prespecified in the analysis plan, all of the above analyses were run with an alternative cutoff for high-pass filtering, i.e. 1 Hz. The results were highly similar to the results obtained for the 0.1 Hz cutoff described above.

## Discussion

In this quality control study, we evaluated the performance of a newly installed OPM-MEG system. Specifically, using an established auditory MMN paradigm, we focused on assessing construct validity (i.e. consistency of MMN results with previously published MMN findings from EEG and SQUID-based MEG) and test-retest reliability (i.e. consistency of MMN amplitude and latency across two separate sessions). Overall, we find that OPM-MEG shows MMN responses that are well compatible with previously reported MMN results with regard to timing and topography. We also find excellent visual consistency of MMN topographies and timeseries across our two sessions. Finally, our results show good test-retest reliability for MMN amplitude at the sensor level, but insufficient reliability for MMN latency.

With regard to construct validity (i.e. comparison with previous MMN results in the EEG/MEG literature), the times of MMN peaks we found with OPM-MEG seem to occur a little earlier than MMN peaks reported in previous EEG (e.g. Weber et al. 2022) or SQUID-based MEG studies (e.g. Maess et al. 2007) but are within the general time window reported for MMN activations. The analysis based on the radial field (z) component allowed for comparing the topography of MMN responses to previous MMN reports based on classical SQUID-based MEG. Here, the study by Maess et al. (2007) provides a good reference point for comparison since it also used a frequency deviant MMN paradigm and shows plots of the activation topography at different time points. Overall, the activation topography we found (Figs. 3 and 4) matches this previous report well.

Concerning out test-retest reliability findings, previous analyses of MMN test-retest reliability using SQUID-based MEG also showed lower reliability for MMN latency than for MMN amplitude (Recasens & Uhlhaas 2017). However, the difference in our case is much larger, and the test-retest results for cluster-level L2 norm based data even indicate a negative ICC value, which is hardly plausible as it implies that within-subject variability is greater than between-subject variability. Furthermore, the poor ICC values for MMN amplitude are difficult to reconcile with the good reproducibility of hemisphere-specific MMN traces across sessions, as suggested by visual inspection of Figures 2 and 4 above. One of the reasons for the low test-retest reliability of latency could be that some participants did not show strong MMN responses, which would result in consistently low amplitude but highly varying latency due to the absence of a clear peak within the time window considered. Such a scenario would not harm the ICC for amplitude but could strongly affect the ICC for latency. Further analyses of subject-level data will be needed to confirm or reject this hypothetical explanation.

In addition to the ongoing investigation of the poor test-retest reliability of MMN latency values in the current analyses, two additional analyses might improve test-retest reliability estimates. First, an obvious next step is to move the analysis to the level of neural sources, following source reconstruction. This choice was made by numerous previous test-retest reliability studies of the MMN, including the most recent MEG study by Recasens & Uhlhaas (2017). Second, it is worth pointing out again that, in this study, we evaluated our OPM-MEG “out of the box”, i.e. directly after installation and without any further optimisation. In particular, one potentially important improvement is the development and deployment of methods for realigning sensor positions across participants. In this study, we only used manual checks of helmet position during data acquisition, but did not employ any post-acquisition methods focal registration. As a consequence, the signals at any given sensor may show variability across participants that is partially due to suboptimal realignment. We expect that addressing this issue will improve our estimates of test-retest reliability. Having said this, given that MMN amplitude showed good test-retest reliability in our current analyses, it is not immediately clear why this potential issue would have preferentially affected MMN latency.

In conclusion, this quality control study of an “out-of-the-box” OPM-MEG system demonstrated good construct validity and mostly good test-retest reliability (with the exception of MMN latency) for one of the most frequently applied paradigms in EEG/MEG research, the auditory MMN. This lends further support to OPM-MEG as an attractive method for human brain activity measurements in routine practice and, more specifically, highlights its suitability for investigations of auditory MMN in health and disease. We are presently in the process of clarifying the reasons behind the surprisingly poor reliability of MMN latency values (which are at odds with the good reliability of amplitude measures and the excellent inter-session group-level reliability of MMN responses) and, in particular, will explore the impact of methods for improved realignment of sensor positions across subjects.

## Acknowledgment

This work was supported by the Swiss National Science Foundation (grant no. 326030_205677), the University of Zurich, ETH Zurich, and the René und Susanne Stiftung via the ETH Foundation.

